# Does data cleaning improve brain state classification?

**DOI:** 10.1101/533075

**Authors:** Steven L. Meisler, Michael J. Kahana, Youssef Ezzyat

## Abstract

**Background:** Neuroscientists routinely seek to identify and remove noisy or artifactual observations from their data. They do so with the belief that removing such data improves power to detect relations between neural activity and behavior, which are often subtle and can be overwhelmed by noise. Whereas standard methods can exclude certain well-defined noise sources (e.g., 50/60 Hz electrical noise), in many situations there is not a clear difference between noise and signals so it is not obvious how to separate the two. Here we ask whether methods routinely used to “clean” human electrophysiological recordings lead to greater power to detect brain-behavior relations.

**New Method:** This, to the authors’ knowledge, is the first large-scale simultaneous evaluation of multiple commonly used methods for removing noise from intracranial EEG recordings.

**Results:** We find that several commonly used data cleaning methods (automated methods based on statistical signal properties and manual methods based on expert review) do not increase the power to detect univariate and multivariate electrophysiological biomarkers of successful episodic memory encoding, a well-characterized broadband pattern of neural activity observed across the brain.

**Comparison with Existing Methods:** Researchers may be more likely to increase statistical power to detect physiological phenomena of interest by allocating resources away from cleaning noisy data and toward collecting more within-patient observations.

**Conclusions:** These findings highlight the challenge of partitioning signal and noise in the analysis of brain-behavior relations, and suggest increasing sample size and numbers of observations, rather than data cleaning, as the best approach to improving statistical power.

## Introduction

All measures of neural activity comprise mixtures of sources, some reflecting the physiological processes we seek to understand and others reflecting physiological or non-physiological sources that contaminate our signals of interest. In human neurophysiological recordings, nuisance signals may reflect electrical potentials produced outside of the brain (e.g., those produced by muscle or eye movements), electrical signals caused by sources within the brain other than those we seek to observe (e.g., epileptic activity contaminating signals related to normal cognition), or even sources outside of the body entirely, such as 50/60 Hz line noise or crosstalk between electrical channels within the recording system. To reduce the influence of these nuisance signals, and thereby increase the statistical power to detect brain-behavior correlations, researchers employ a variety of data cleaning methods. Although the use of data cleaning methods is nearly ubiquitous in neuroscientific research, it remains unknown whether these methods actually increase the power to detect relations between physiology and behavior. The goal of this report is to systematically investigate the effectiveness of commonly used data cleaning approaches.

As an example case, we surveyed the literature on intracranial electroencephalography (iEEG) studies of human episodic memory encoding, a fairly restrictive domain. Electrophysiological studies of memory encoding have consistently demonstrated that increases in broadband high-frequency activity, a correlate of multi-unit neural activity (Rutishauser, Ross, Mamelak, & Schuman, 2010; Manning, Jacobs, Fried, & Kahana, 2009; Winawer et al., 2013), during encoding predicts subsequent recall, an effect seen across a broad network of brain regions including hippocampus, medial temporal lobe (MTL), and lateral prefrontal cortex (Burke et al., 2014; Long, Burke, & Kahana, 2014; Long & Kahana, 2015; Sederberg et al., 2006). We used this high-frequency signal in our univariate analyses described below because increased high-frequency activity is also linked to successful cognitive performance across a range of domains (Crone, Sinai, & Korzeniewska, 2006; Cheyne, Bells, Ferrari, Gaetz, & Bostan, 2008; K. J. Miller et al., 2007; Hermes, Miller, Wandell, & Winawer, 2015; Chang et al., 2011). These broadly distributed high-frequency patterns have also been combined with low-frequency signals into novel multivariate models for guiding closed-loop systems to detect memory encoding states (Ezzyat et al., 2018; Ezzyat & Rizzuto, 2018). We therefore applied our analyses to both a univariate and multivariate measure of memory encoding, in order to determine whether data cleaning would differentially impact the two approaches.

In our survey of the iEEG episodic memory literature, we identified six commonly used procedures aimed at removing or attenuating putative noise sources prior to hypothesis testing (see Table 1). These six methods fell broadly into three categories: (1) methods based on statistical properties of the voltage timeseries (variance and kurtosis); (2) manual annotation of data based on suspected epileptic activity over channels or epochs; and (3) common average vs. bipolar referencing. Our goal was to evaluate the effectiveness of each approach in increasing the power to detect electrophysiological correlates of memory encoding. We applied each of these methods separately at both the channel and epoch levels, and for the automated techniques we compared the effect of removing putatively noisy data with quantity-matched random data removal. We applied the methods to a dataset of 127 patient participants (although not all patients had manual annotations from expert review), each contributing hundreds of observations during a memory encoding task. Taking advantage of the large quantity of within-subject data, we further evaluated each data cleaning method as a function of the quantity of data collected (# of channels and # of events) to determine whether the data quantity interacted with a particular method’s effectiveness in increasing statistical power.

**Table 1:** Preprocessing approaches used in prior iEEG studies of human episodic memory.

|  | METHODS |  |  |  |
| --- | --- | --- | --- | --- |
|  | Bipolar | Kurtosis | Standard Deviation | Epileptic Activity |
| Raghavachari et al., 2001 |  |  |  | X |
| Caplan et al., 2001 |  |  |  | X |
| Sederberg et al., 2003 |  | X |  |  |
| Ekstrom et al., 2005 |  |  |  | X |
| Greco et al., 2007 |  | X |  |  |
| Sederberg et al., 2007 |  | X |  |  |
| van Vugt et al., 2009 |  | X |  |  |
| van Vugt et al., 2010 |  | X |  |  |
| Nolan et al., 2010 |  |  | X |  |
| Whitmer et al., 2010 |  |  |  |  |
| Dastjerdi et al., 2011 |  |  |  | X |
| Burke et al., 2013 | X |  |  |  |
| Swann et al., 2013 |  |  |  | X |
| Burke et al., 2014 | X |  |  |  |
| Lega et al., 2014 |  | X |  |  |
| Rangarajan et al., 2014 |  |  | X | X |
| Zavala et al., 2014 | X |  |  |  |
| Greenberg et al., 2015 | X | X |  |  |
| Zhang and Jacobs, 2015 |  | X |  |  |
| Haque et al., 2015 | X |  |  |  |
| Kim et al., 2016 |  |  |  | X |
| Vass et al., 2016 |  |  |  | X |
| Piai et al., 2016 | X |  |  |  |
| Fonken et al., 2016 |  |  | X | X |
| García-Cordero et al., 2017 |  |  |  | X |
| Sheehan et al., 2018 | X | X | X |  |

Because there is no widely adopted standard for cleaning noise from iEEG data, investigators are left to apply methods which they have used in the past or those used by others in the field without knowing which methods are likely to be effective. For methods based on statistical properties of the timeseries, which can be scripted and applied automatically, it is particularly easy for researchers to (consciously or not) evaluate multiple thresholds and methods and select the one that leads to the “cleanest” end result (Simmons, Nelson, & Simonsohn, 2011; Silberzahn et al., 2018). In the case of more labor intensive methods like identifying epileptic artifacts that require human annotation, subjectivity in criterion setting means that the rejected data will vary from one expert annotator to the next. Manual methods also have the obvious downside of requiring more time to carry out; to the extent that manual methods are effective at identifying and removing noise, this may be worth the investment of time and effort. However if manual annotation does not, on average, have any benefit then it simply squanders resources that could be better allocated elsewhere.

Across univariate and multivariate measures we find that the majority of commonly used approaches to identifying and removing noisy epochs or channels do not increase statistical power. For methods based on using a numeric threshold on statistical measures of the voltage timeseries (e.g. signal variance or kurtosis), typically-used liberal criteria (*z* > 2.5) do not increase statistical power compared to randomly removing an equivalent number of epochs or channels, while, as expected, using more conservative thresholds that exclude more data lead to reductions in statistical power. For methods based on using human manual annotation to identify epileptic channels or epochs, the overall effect is similar in that removing noisy data does not increase statistical power. We investigated whether the statistical or manual approaches might be more effective when applied to smaller datasets, however a within-subject parametric analysis indicated this was not the case. The major exception to this pattern of null results occurred when comparing bipolar and common average referencing approaches, where bipolar referencing significantly outperformed common average referencing in several analyses, particularly for the multivariate paradigm. The data show that the choice of referencing has a profound impact on iEEG analysis broadband measures of neural activity, while other commonly used noise removal approaches do not, and suggest that researchers will benefit from allocating resources away from post-processing noise and toward collecting more within-subject data.

## Materials and Methods

### Participants

We analyzed data from 127 neurosurgical patients with medication-resistant epilepsy who participated in at least three sessions of a verbal free recall memory task (described below) as part of an ongoing research collaboration coordinated by the University of Pennsylvania. As part of their clinical workup, subjects had electrodes implanted intracranially, either on the cortical surface or within the brain parenchyma–recordings from these electrodes during performance of the memory task were then used in the analyses described in this report. Subjects performed the memory task using a laptop at the bedside, and were tested at one of the following centers: Thomas Jefferson University Hospital (Philadelphia, PA), University of Texas Southwestern Medical Center (Dallas, TX), Emory University Hospital (Atlanta, GA), Dartmouth-Hitchcock Medical Center (Lebanon, NH), Hospital of the University of Pennsylvania (Philadelphia, PA), Mayo Clinic (Rochester, MN), and Columbia University Medical Center (New York, NY). The Institutional Review Board at each center approved the research protocol, and informed consent was obtained from each participant.

### Free Recall Task

In a free recall task, a subject is presented with a list of words that he or she later tries to recall in any order (Glanzer, 1969). During the encoding phase, 12 nouns (http://memory.psych.upenn.edu/WordPools) were presented one at a time on a computer screen in the subject’s primary language (either English or Spanish). Each word remained on screen for 1600 milliseconds, followed by a blank inter-stimulus interval of between 750-1000 milliseconds (randomly drawn from a uniform distribution). Following the final word in the list, there was a 20 second distractor period, during which the subject used the laptop keyboard to answer arithmetic problems of the form A + B + C = ?, where A, B, and C were random integers between 0 and 9. After the delay ended, an 800 Hz auditory tone was played and a row of asterisks was displayed on the screen (simultaneously for 300 milliseconds), which signaled the beginning of the recall period (30 s). Subjects were instructed to recall as many words from the preceding list as could be remembered, in any order. Figure 1 illustrates the design of the task. Subjects completed up to 25 lists in a single session. Audio responses from the sessions were digitally recorded and manually parsed and annotated offline using Penn TotalRecall (http://memory.psych.upenn.edu/TotalRecall). There were two versions of the task: in one version the lists of words were constructed to minimize semantic relationships between the words; in the other version, four words from each of three semantic categories (e.g. DESK, CHAIR, TABLE, SOFA) were presented within the same list. Subjects could participate in one or both tasks across different sessions, however within a given session, only one task was used.

**Figure 1:**
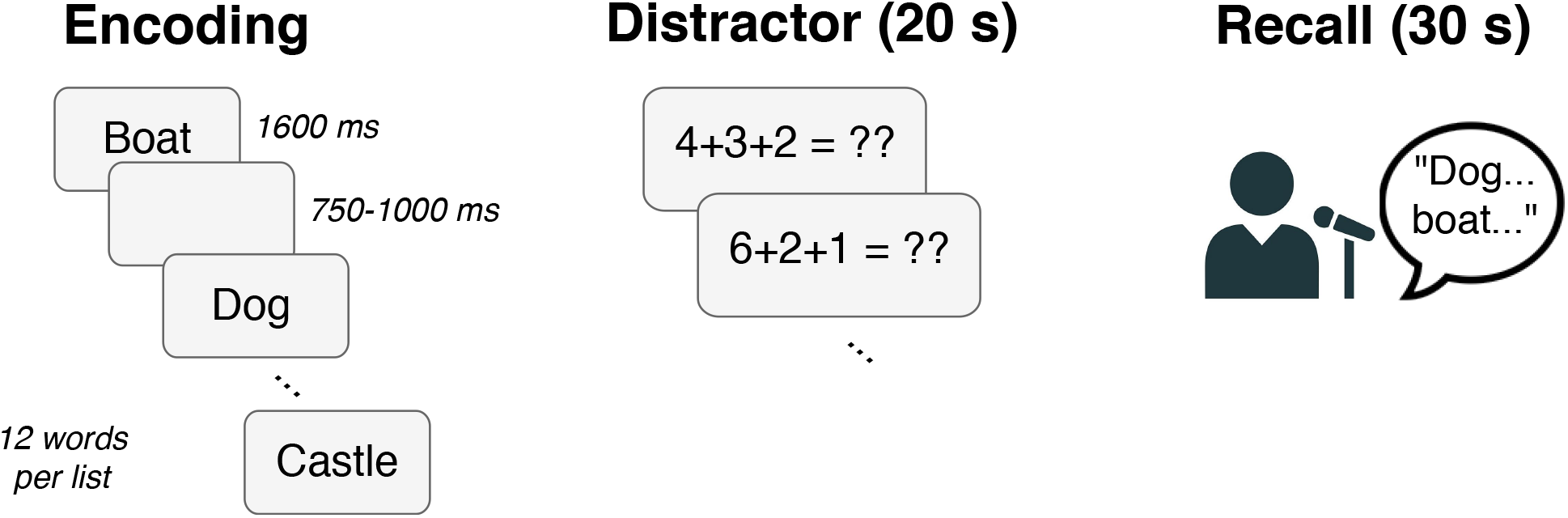
Schematic diagram of a single list presentation during the free recall task.

### Electrophysiological Recordings

iEEG data were gathered from both depth and subdural cortical surface electrodes. Subdural cortical surface leads were arranged in both strip and grid configurations. The types of electrodes used for recording varied across institutions based on clinician preference. Across the sites, the following electrode models were employed: PMT Depthalon (0.8 mm), AdTech Spencer RD (0.86 mm), AdTech Spencer SD (1.12 mm), AdTech Behnke-Fried (1.28 mm), and AdTech subdural strips and grids (2.3 mm).

iEEG data were recorded with one of the following clinical EEG interfaces, depending on institutional preference: Nihon Kohden EEG-1200, Natus XLTek EMU 128 or Grass Aura-LTM64. Electrode locations varied across subjects and were determined strictly by clinical monitoring needs. Depending on the amplifier and the preference of the clinical team, the signals were sampled at either 500, 1000, or 1600 Hz and were referenced to a common contact placed intracranially, on the scalp, or on the mastoid process. We notch filtered (Butterworth, 4th order, frequency range = center ±2 Hz) all timeseries at 60, 120, and 180 Hz to reduce line noise.

### Bipolar vs. Common Average Referencing

Bipolar referencing involves referencing pairs of neighboring electrodes against each other, as opposed to common average referencing, which references against the average signal across electrodes (Nunez & Srinivasan, 2006). Bipolar referencing is thought to minimize the impact of reference channel noise on other channels (Burke et al., 2013), however a large-scale direct comparison of bipolar and common average reference schemes using a combination of subdural and depth electrodes has not been reported. We therefore compared bipolar and common average referencing to answer the question of whether there was any benefit, in terms of increasing statistical power by attenuating noise, to using bipolar referencing over common average referencing. Bipolar pairs were created for every group of two adjacent contacts on every depth, strip, and grid in a patient’s montage. The timeseries for each bipolar pair is the difference between the signals in each electrode. We use the term *channel* in this paper to denote either a common average referenced signal from a single electrode, or a bipolar referenced signal from two neighboring electrodes, depending on the modality of the data of interest.

### General Procedure

The goal of this study was to use a large dataset to evaluate the effects of several methods for identifying and removing noise in iEEG recordings. We measured the effect of each method by determining the impact on the subsequent memory effect (SME), a well-characterized biomarker of successful memory encoding, during a free recall memory task. As described below, we separately assessed the effect on a univariate and multivariate measure of the SME, in order to answer the question of whether data removal had a differential impact on statistical power for uni- and multivariate measures. For both the univariate and multivariate analyses of the SME, we used spectral decomposition of the iEEG recordings to identify power in specific frequency bands; we describe the details in the next section. A schematic detailing our procedures for statistical thresholding and computing classifier metrics can be found in Figure S1.

### Spectral Decomposition

We extracted all word encoding intervals from each subject’s recordings (1600 ms following each word onset) and used Morelet wavelet convolution (wave number = 5) to spectrally decompose each epoch. We included a 1500 ms buffer period before and after each word encoding epoch before convolution to mitigate edge artifacts, and subsequently discarded the buffer period before further analysis.

For the multivariate analysis, we used eight wavelets with center frequencies logarithmically-spaced between 3 and 175 Hz (3.0, 5.4, 9.6, 17.1, 30.6, 54.8, 97.9, 175.0). For the univariate analysis, we also used eight wavelets, but focused on the high frequency range (between 70 and 200 Hz; 70.0, 81.3, 94.5, 109.8, 127.5, 148.2, 172.1, 200.0). We then log-transformed and averaged the resulting powers over the 1600 ms word encoding interval. We then normalized the powers (*z*-transform) across word encoding epochs, separately within each session, channel and frequency (where relevant, the *z*-transformation was applied after applying one of the evaluated noise attenuation methods).

### Multivariate Classification

The goal of the multivariate analysis was to use patterns of spectral power (across frequencies and channels) during encoding to classify individual word encoding epochs as recalled/not recalled. We used a logistic regression classifier implemented with Python’s *scikit-learn* (Pedregosa et al., 2011) library, and using a balanced class weight to account for the difference in the base rate of recalled/not recalled trials. To reduce overfitting, we used L2 penalization (Hastie, Tibshirani, & Friedman, 2001) with penalty parameter *C* = 2.4 × 10^−4^, following protocols from previous free recall studies with a similar subject cohort (Ezzyat et al., 2017). We trained the classifier on feature matrices comprised of the *z*-scored powers (the input) and recall outcomes (binary -”recalled” or “not recalled”; the output). We used *N* − 1 cross-validation at the list-level as follows: we trained the model using all epochs with the exception of those corresponding to a single held-out list (12 epochs); we then tested the trained model on the held-out list to generate predicted probabilities of recall for each held-out epoch. We then iterated this procedure until all lists had been held-out once. We created receiver operating characteristic (ROC) curves for each subject using the cross-validated classifier probabilities and the true recalled/not recalled labels. An ROC curve plots the true positive rate against the false positive rate at various probability thresholds. We assessed model performance using area under the ROC curve (AUC), with an AUC of 0.5 representing chance (Hanley & McNeil, 1982).

### Univariate Measure of SME

The goal of the univariate analyses was to compare spectral power in the high-frequency range (70–200 Hz) during the encoding period of recalled vs. not recalled words. Univariate high-frequency power during memory encoding across the brain is well-known to differentiate subsequently recalled and not recalled words (Burke et al., 2014). We averaged the *z*-scored powers over the word encoding period (0–1600 ms) separately for each epoch × frequency bin × channel, and used Welch’s *t*-test *across* epochs *within* each channel to compare power for subsequently recalled vs. not recalled epochs (we employed Welch’s test due to the possibility for different *N*s and variances for the recalled/not recalled distributions). We then averaged the *t*-statistic across all channels to derive a summary measure of the univariate subsequent memory analysis.

### Channel and Epoch-Based Statistical Thresholding

We evaluated the use of standard deviation and kurtosis thresholds for excluding channels from analysis. After collecting the timeseries data and separating them into epochs, we calculated the standard deviation and kurtosis of each epoch on each channel. Within each channel, we then computed the average standard deviation and average kurtosis across epochs. Then, we *z*-scored these metrics separately across channels, and we evaluated the effect of excluding channels on the basis of standard deviation/kurtosis values at various thresholds in the distribution across channels for a given subject.

We excluded a channel if its kurtosis or standard deviation *z*-score exceeded a given threshold. We evaluated the effect of thresholds at intervals of 0.5 between and including 0 and 3. As a benchmark for comparison, we also compared the threshold-based approach to randomly removing an equivalent number of channels at each threshold. We repeated the random removal 50 times, and the reported AUC/*t*-statistic for random removal represents the average across iterations.

Similar to the analysis described in the previous paragraphs, we also tested the effect of excluding individual epochs based on their kurtosis and standard deviation. Within each epoch, we averaged the metric of interest (standard deviation, kurtosis) across channels. We proceeded to *z*-score these averages across epochs. We randomly removed epochs similarly to the channel randomization in the previous section. This process is also repeated 50 times, with the reported random AUC / *t*-statistic being the average of the 50 iterations. The multi- and univariate results can be found in figures 2 and 3, respectively. While the previous sections described automated statistical thresholding, the next two sections detail removing data based on manual/semi-manual physician identification.

**Figure 2:**
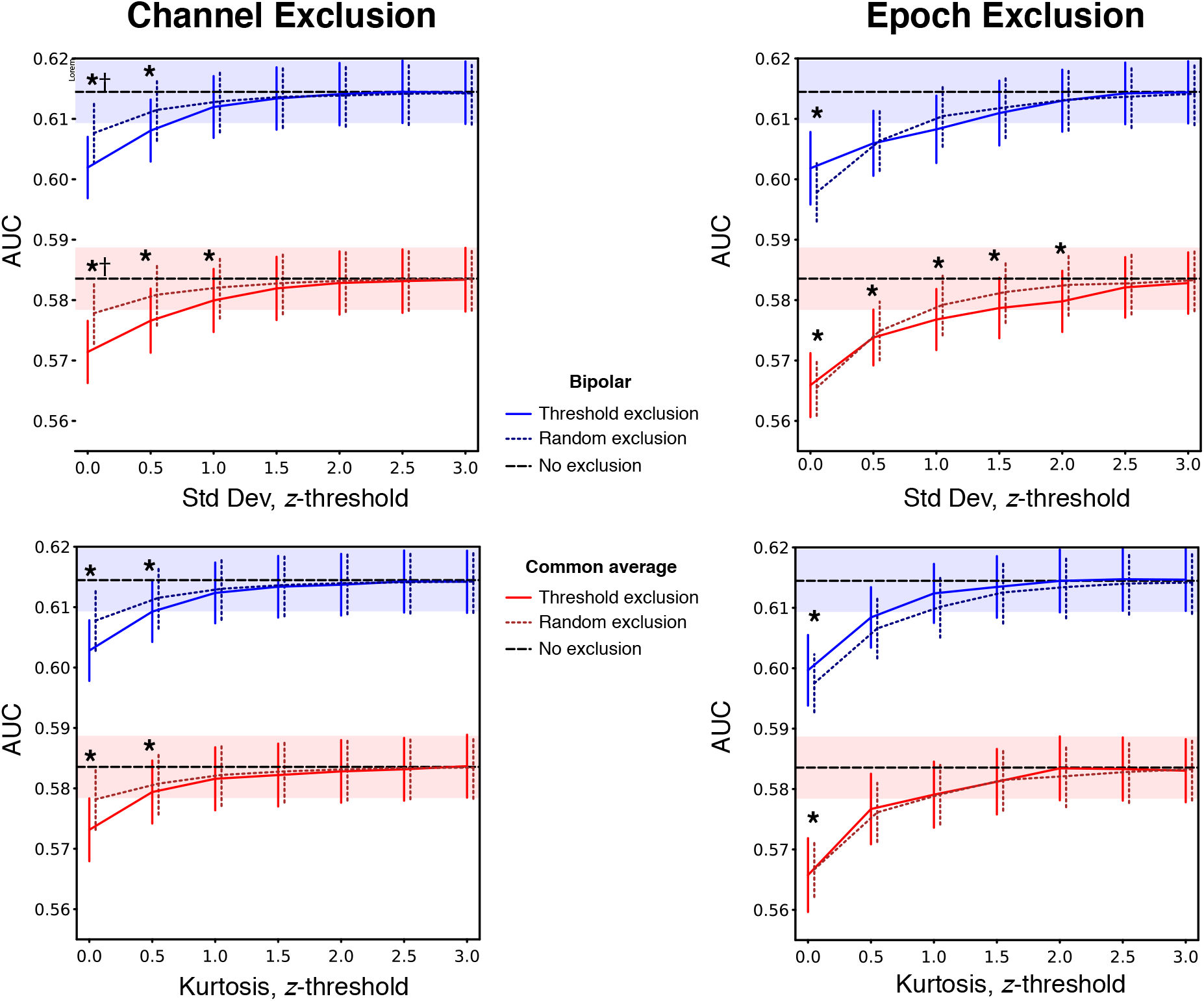
Statistical thresholding applied to the full multivariate dataset. The columns are differentiated by the type of data excluded (channels vs. epochs), and the rows are differentiated by the basis of exclusion (standard deviation vs. kurtosis). Referencing scheme (common average vs. bipolar) is indicated by color in all figure panels. The black dashed lines represent the respective values for the *No exclusion* baseline. * Indicates FDR-corrected *q* < 0.05 compared to *No exclusion* baseline. † Denotes FDR-corrected *q* < 0.05 compared to *Random exclusion*. The shaded region surrounding those lines represent the SEM. Lines are horizontally offset for visual clarity. Error bars represent SEM.

**Figure 3:**
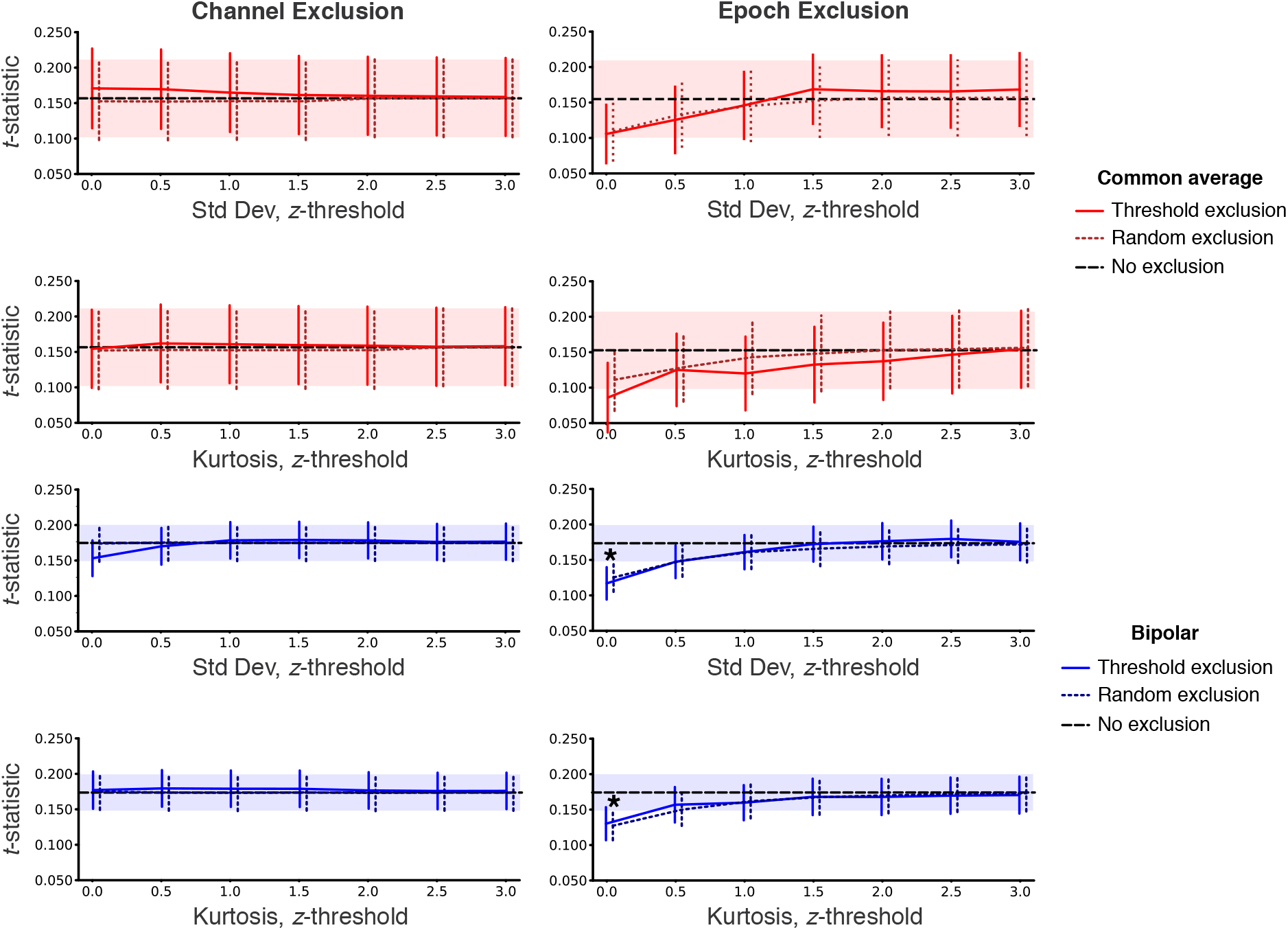
Statistical thresholding applied to the full univariate dataset. The columns are differentiated by the type of data excluded (channels vs. epochs), and the rows are differentiated by the basis of exclusion (standard deviation vs. kurtosis). Referencing scheme (common average vs. bipolar) is indicated by color in all figure panels. The black dashed lines represent the respective values for the *No exclusion* baseline. * Indicates FDR-corrected *q* < 0.05 compared to baseline. The shaded region surrounding those lines represent the SEM. Lines are horizontally offset for visual clarity. Error bars represent SEM.

### Neurologist-Based Channel Exclusion

At each participating medical center, clinicians identified electrodes in regions associated with three types of abnormal neural activities: interictal spiking, seizure onsets, and lesions. Electrode indications were stored in a subject-specific text file, although one did not exist for every subject in this experimental cohort. For the 94 subjects who had these annotations, the information was parsed to identify common average referenced channels associated with the abnormal categories. Any bipolar referenced channel that included at least one such electrode was also grouped as belonging to the abnormal category associated with the channel(s). All possible combinations of categories were removed before performing spectral decomposition and classification. The resulting AUCs / *t*-statistics may be found in Figure 4.

**Figure 4:**
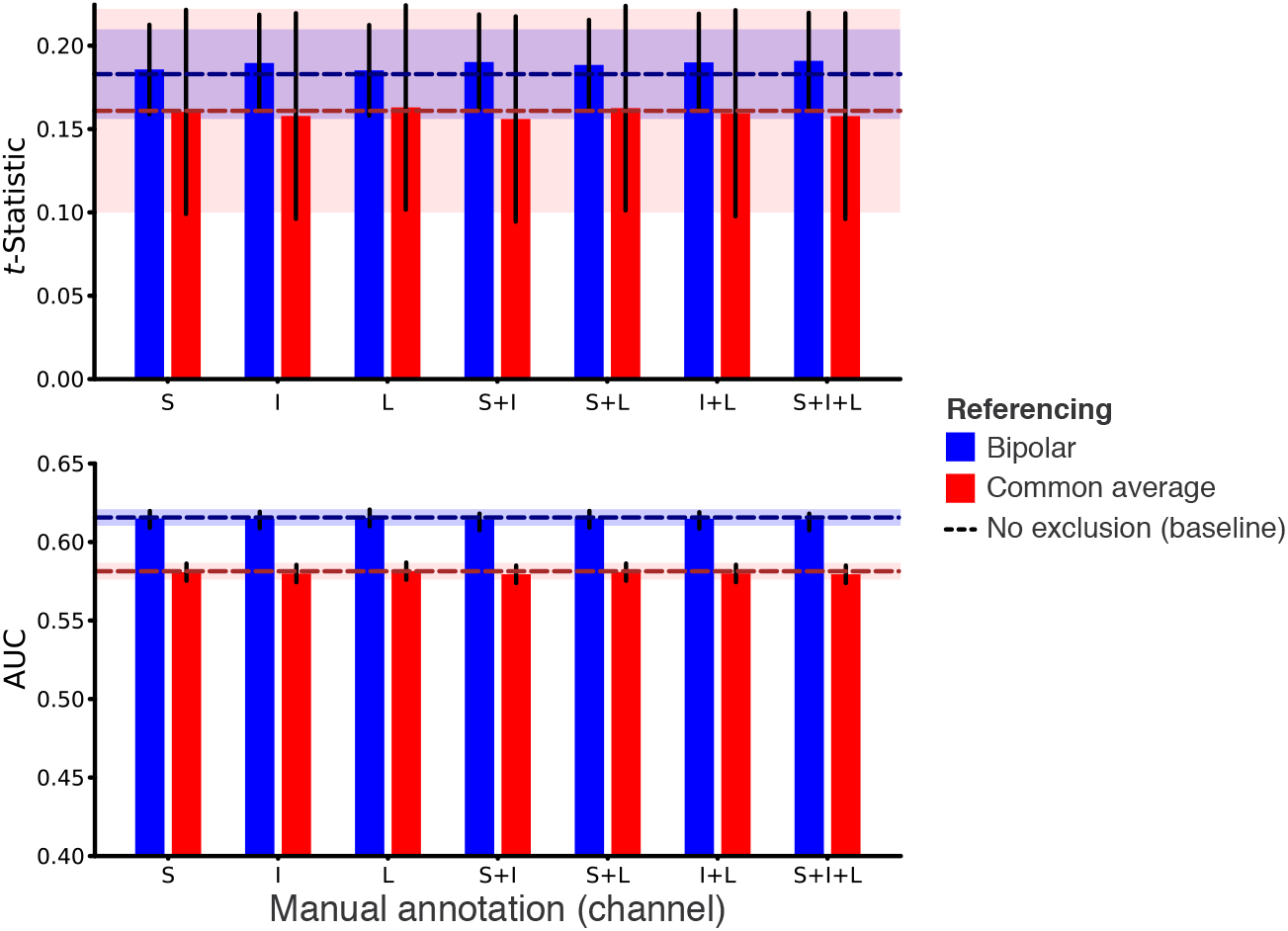
Manual channel exclusion applied to the full dataset in 94 subjects. Error bars represent SEM. Horizontal dashed lines indicate the *t*-statistic/AUC measure with no data exclusion. Shaded regions indicate standard error of the *No exclusion* baseline conditions. S: Seizure onset, I: Interictal spiking, L: Lesion. For all combinations of data exclusion there was no difference in comparison with the no exclusion baseline.

### Neurologist-Based Epoch Exclusion

Recently, a semi-automatic method has been evaluated to identify epochs with high frequency oscillatory (HFO) activity, a characteristic of epileptic EEG and iEEG (Waldman et al., 2018; Shimamoto et al., 2018; Weiss et al., 2018). The algorithm, run only on depth electrodes for optimal signal-to-noise, annotates an epoch with one of the following labels that are then visually validated: 0) No HFO event detected, 1) Ripple superimposed in an interictal discharge, 2) Sharply contoured epileptiform spike, 3) Ripple on oscillation, 4) Fast ripple superimposed in an interictal discharge, 5) Fast ripple on oscillation. Following the guidance of the developers of the method, we removed as artifacts any epoch with an annotation of 1, 2, and/or 4. Then spectral decomposition and *t*-stat and AUC calculations proceeded similarly to other analyses, with the exception that only depth electrodes were used to build the classifiers. Of the original subject cohort, 28 patients had been analyzed with this technique and had HFO epochs to remove. Results for this analysis may be found in Figure 5.

**Figure 5:**
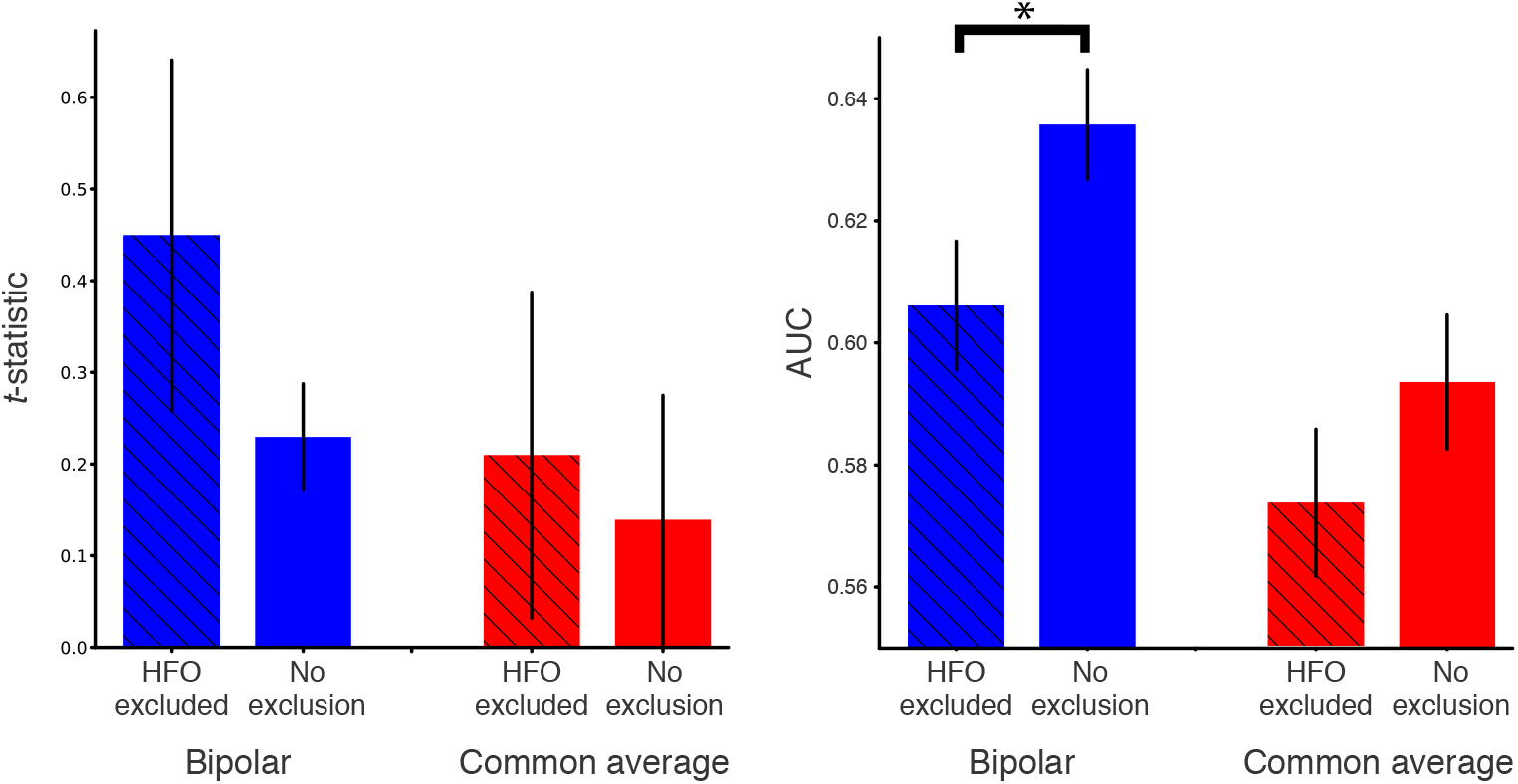
Neurologist-identified epoch exclusion applied to depth electrodes in 28 subjects. Hatched bars represent outcome metrics after HFO epochs were excluded, and solid bars represent baseline. * Denotes significant difference from the *No exclusion* baseline (Wilcoxon signed-rank test; FDR-corrected *q* = 0.01).

### Data Subsampling

In addition to evaluating the effect of applying the preceding methods to a subject’s full dataset, we were also interested in how noise removal methods might differ in their effects on large and small within-subject datasets. To address this question, for all previously described methods, we repeated the analyses using variously-sized subsets of data within subject. We randomly sampled 20, 40, 60, or 80% of either (1) each subject’s full set of channels or (2) each subject’s full set of epoch lists. This process was iterated 50 times per session and sampling percentage, and the reported result for the subject is the average across iterations and sessions. For methods that involved statistical thresholding, *z*-score thresholds below 2 were not tested since *post-hoc* analysis of the full datasets showed they did not improve classifier performance. Subsampling results may be found in the Supplementary Materials.

### Statistics

All results are presented as the mean ± standard error of the mean (SEM) unless noted otherwise. Any horizontal dashed lines in results figures represent baseline metrics present when no data are removed. The shaded region around them represents the SEM surrounding the baselines. All statistical tests, with the exception of those in Figure 5, were multiple comparisons corrected using false discovery rate (?, ?). For the analysis in Figure 5, because there were many fewer tests, we used Bonferroni correction.

### Data Availability

Electrophysiological data may be found at http://memory.psych.upenn.edu/Electrophysiological_Data. The Python Time Series Analysis (PTSA) toolbox (https://github.com/pennmem/ptsa_new) was used to analyze the data. Custom Python scripts are available upon request from Steven Meisler.

## Results

Figures 2 – 5 show the impact of the cleaning methods on classification outcomes.

### Channel and Epoch-Based Statistical Thresholding

As shown in Figure 2, removing data based on their variance or kurtosis did not increase multivariate classifier performance above baseline. As denoted by asterisks in the figure, the random removal (dotted lines) occasionally performed significantly better than the true exclusion (Wilcoxon signed-rank test; FDR-corrected *q* < 0.05). At every threshold, the AUCs for bipolar pairs were significantly greater than their common average counterparts (Wilcoxon signed-rank test; *p* < 1 × 10^−10^). Similarly, in the univariate measure, depicted in Figure 3, removing data at any threshold never led to a significant increase in *t* above the *No exclusion* condition.

We also assessed the performance of noise removal methods for multivariate and univariate measures, as a function of the amount of within-subject data available. We subsampled each participant’s dataset by either channels or epochs and performed the multivariate and univariate analyses for the subsampled data. As was the case for the full dataset, automated noise removal methods did not increase the size of the subsequent memory effect in smaller datasets (Fig S6) and S7).

### Manual Channel Exclusion

*N* = 94 subjects had manually-identified channels to exclude. Removing channels associated with abnormal neural activity did not significantly improve classifier performance. As depicted in Figure 4, AUCs after removal never exceed the *No exclusion* baseline. Similar to the previous results, the bipolar AUCs significantly exceed those of common average data (Wilcoxon signed-rank test; *p* < 1 × 10^−10^).

### Neurologist-Based Epoch Exclusion

*N* = 28 subjects had annotated HFO epochs to exclude. As depicted in Figure 5, the average *t*-statistic for the data after HFO-exclusion was higher than baseline in both modalities, although not significantly. The opposite effect was observed in the multivariate results, especially in the bipolar data, in which baseline significantly exceeded AUCs after HFO removal (Wilcoxon signed-rank test; FDR-corrected *q* = 0.01)

## Discussion

Neuroscientists regularly identify and exclude data through both automatic and manual means, with the underlying assumption that doing so will increase the statistical power for detecting physiological phenomena. In this study, we systematically evaluated the impact of several preprocessing methods commonly employed in iEEG studies of human memory with the goal of determining if the approaches are effective at increasing statistical power at both a univariate and multivariate level. The methods we analyzed included bipolar referencing as well as removing both channels and epochs through automatic statistical thresholding and manual identification. In our cohort of 127 subjects, we also tested the efficacy of preprocessing while varying the amount of within-patient data, both in terms of the number of electrodes and session length. We found that the preprocessing methods that involved data (e.g. epoch and channel) exclusion, did not increase the statistical reliability of the subsequent memory effect. In some cases, primarily in the multivariate paradigm, the targeted data removal decreased statistical power more than random exclusion did. The major exception to this general finding was using bipolar referencing. Finally, when varying the amount of within-patient data through subsampling, we did not find an interaction between the effectiveness of noise exclusion and the size of the underlying dataset.

Although our results are consistent with the interpretation that commonly used data cleaning methods do not increase statistical power, these methods may nonetheless be valuable in situations where the artifact or noise can be well characterized, and/or is correlated with the behavior of interest. For example, one could imagine a scenario in which confounding electromyogram (EMG) or electrooculogram (EOG) activity from muscle contractions (e.g. due to nervousness or concentration, or eyeblinks) could be correlated with memory performance. In such a situation, a researcher would certainly want to eliminate contamination from such artifacts in order to maximize the purity of the signal that actually comes from the brain. However, being able to do this relies on the assumption that a researcher has a good model of the artifact and has strong evidence that the artifact is correlated with the behavior or neurophysiological signal of interest. If the artifact is not strongly related to the behavior, and if the ability to distinguish artifact from signal is poor, then removing artifacts is more likely to remove a mixture of valid and artifactual data, than strictly artifactual data. This can complicate conclusions drawn from the “cleaned” data will be representative of the true relationship between physiology and behavior. In the case of bipolar recordings from intracranially implanted electrodes, one would not expect to see significant EMG and EOG artifacts. In situations in which the characteristics of the artifact are known (e.g. 50/60 Hz line noise), data cleaning is advantageous. However, our results suggest that for cases in which the distinction between signal and noise is less obvious, the researcher will be better off collecting and analyzing as much data as possible.

A second reason to reconsider data cleaning is because it offers researchers additional parameters with which to tune analyses. Because there is no widely accepted approach for intracranial studies of human memory that researchers can adopt *a priori*, there is a greater opportunity for a study to lead to distorted inferences about brain-behavior relationships that will not generalize. Opportunities to collect data from iEEG electrodes are usually scarce (Parvizi & Kastner, 2018), making the recordings themselves highly valuable–ideally researchers would make use of as much of the collected data as possible. Our results suggest that investigators should be empowered to analyze as much of their data as possible, and that a fruitful use of resources would be to collect large numbers of within-subject observations.

It is interesting that targeted data removal sometimes led to smaller estimates of the SME, in particular when using a multivariate measure of the SME, as in Fig. 5. This finding suggests that epilepsy-related HFO events may correlate with memory processes. If this is the case then removing such epochs from the data will be likely to remove informative physiological signal along with the HFO artifact, which would be expected to reduce parameter estimates of the SME. Figure 5 also suggests that multivariate classifiers can make use of memory-related information that may be present in these HFO epochs. This would be consistent with other work in epileptic patients that has shown a relation between epileptic discharges and episodic memory performance (Horak et al., 2017).

Our analyses focused on evaluating the types of data cleaning methods that have been employed in a restricted domain of intracranial EEG research, namely episodic memory. These data cleaning techniques have often been used in conjunction with time-frequency analyses in which voltage timeseries are transformed in order to extract time-varying spectral content. Our results are therefore likely to generalize most readily to other intracranial EEG work in domains that also adopt a spectral decomposition approach (Lachaux, Axmacher, Mormann, Halgren, & Crone, 2012; K. Miller et al., 2007; Voytek et al., 2010). There is also widespread evidence that changes in high-frequency power correlate with cognition (Crone et al., 2006; Cheyne et al., 2008; K. J. Miller et al., 2007; Hermes et al., 2015; Chang et al., 2011), suggesting applications of this work to other domains outside of episodic memory. On the other hand, we believe our data are also important more broadly for researchers employing other methods that emphasize time-domain data representations (Phan, Wachter, & Kahana, Submitted). While the efficacy of particular data cleaning methods will depend on the chosen analysis approach, our data question the broader assumption that reducing the sample size of a dataset by excluding putatively noisy observations leads to increased statistical power.

Our data contribute to a growing literature on optimal methods for referencing electrophysiological data. For applications where the features of interest are power at a priori defined frequencies of interest, future work could evaluate the efficacy of methods for separating the raw data into signal and noise components using eigenvalue decomposition (Nikulin, Nolte, & Curio, 2011), instead of bipolar referencing followed by wavelet convolution. Other data-driven approaches such as independent component analysis (ICA) have also been shown to outperform bipolar referencing in simulations involving relatively small numbers of channels (Michelmann et al., 2018), suggesting an extension of our work that would compare bipolar and ICA-based referencing on a large dataset collected from human participants. Other transformations such as the Laplacian (Li et al., 2018) may help eliminate confounding effects of white matter electrodes in other referencing schemes (Mercier et al., 2017), thereby outperforming a standard bipolar montage. Although a comparison between the bipolar approach and these other methods is beyond the scope of this report, future work should focus on comparing these different approaches.

Generally, whether it be through targeted preprocessing, random exclusion, or subsampling, removing epochs tended to have a larger impact on statistical power than removing electrodes, and this difference was more pronounced in the univariate analysis. The reported *t*-stat was the average of tests across individual electrodes, so removing electrodes did not change the statistical power of the tests, which is determined by the number of epochs. Surprisingly, although removing “noisy” data failed to improve statistical power, bipolar referencing increased statistical power. Although the improvement was modest, it was statistically significant in the multivariate analyses. This effect could be due to the utility of bipolar referencing in minimizing propagation of artifacts on reference channels. This also suggests that local as compared with global spectral changes better characterize variability in memory encoding, at least at high frequencies (Solomon et al., 2017).

Another way to view our results would be that the classifiers, particularly the multivariate logistic regression, were robust to the noise that data exclusion targeted (Tomioka, Aihara, & Müller, 2007). This is consistent with previous work showing that L2-penalized logistic regression classifiers generalized to situations in which electrodes were excluded (Hammon & de Sa, 2007). Logistic regression also proved to be reliable in single-trial EEG classification (Tomioka et al., 2007) and cursor movement (Penny, Roberts, Curran, & Stokes, 2000) for a BCI. Similar robustness has also been observed for classifiers in other domains (Ryali, Supekar, Abrams, & Menon, 2010; Bootkrajang & Kabán, 2012; Carroll & Pederson, 1993; Komarek & Moore, 2003).

## Conclusion

This study tested the impact of various preprocessing methods on multivariate and univariate iEEG measures of episodic memory encoding. Across all tests that involved removing data based on automatic statistical thresholding or manual identification, exclusion did not improved classifier performance. Bipolar referencing, however, improved classifier metrics, particularly in the multivariate paradigm. These results were consistent across varying amounts of within-patient data. The study suggests that many commonly used approaches for removing iEEG noise are likely to be less effective in increasing statistical power than either bipolar referencing or increasing the number of within-participant observations.

## Conflicts of Interest

Dr. Michael Kahana has co-founded a company, Nia Therapeutics, LLC (“Nia”), intended to develop and commercialize brain stimulation therapies for memory restoration. He holds more than 5% equity interest in Nia. The views, opinions, and/or findings contained in this material are those of the authors and should not be interpreted as representing the official views or policies of the Department of Defense or the U.S. Government.

## Acknowledgments

The authors would like to acknowledge Blackrock Microsystems’ generosity in providing neural recording equipment. This work was supported by the DARPA Restoring Active Memory (RAM) program (Cooperative Agreement N66001-14-2-4032), as well as National Institutes of Health grant MH55687 and T32NS091006. This study would not be possible without all of the patients who volunteered their time to participate in the study.

## Supplemental Information

**Figure S1:**
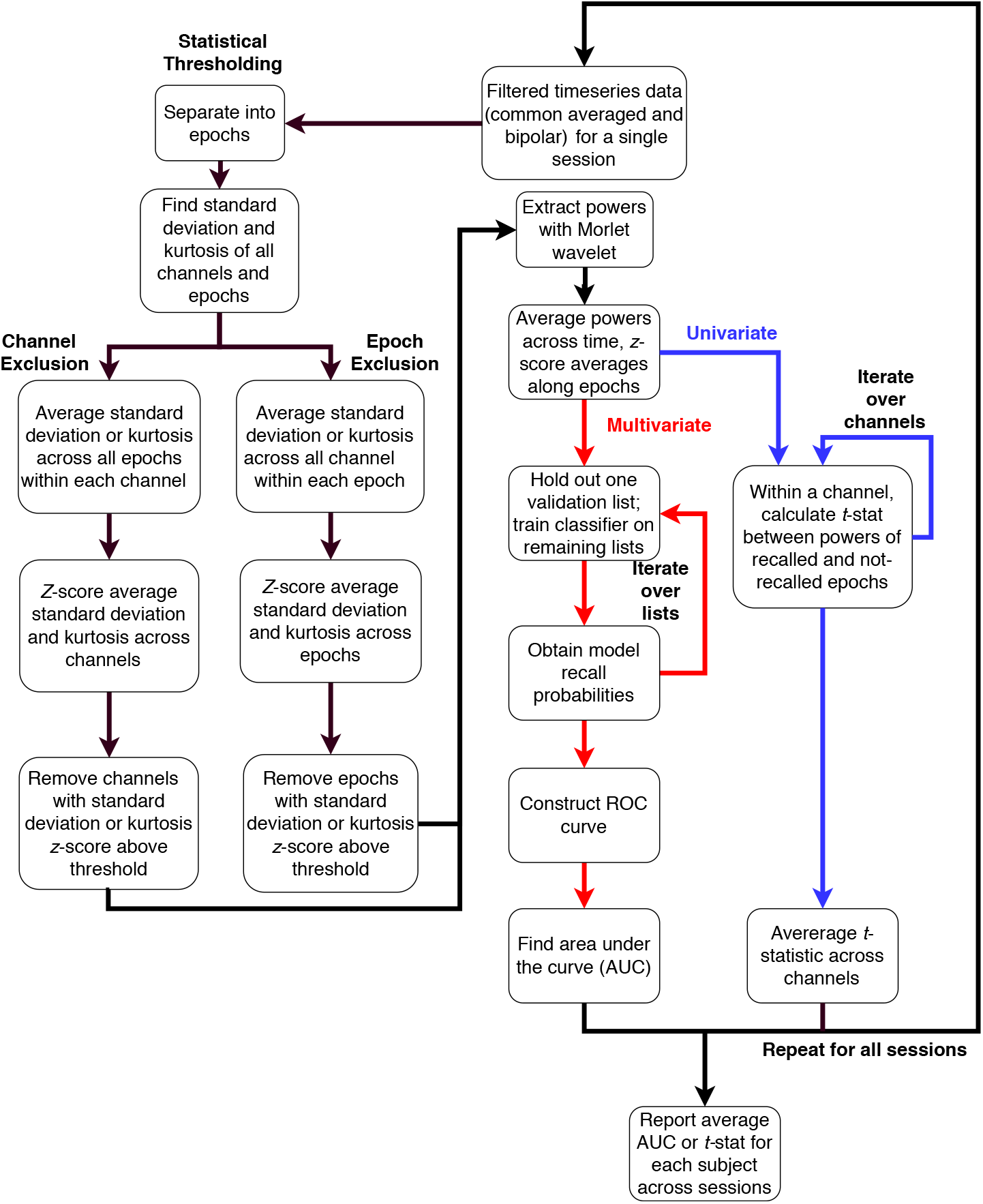
Schematic diagram depicting the statistical thresholding methods in this study. The left side of the diagram depicts how we excluded channels and epochs, and the right side details how we computed the univariate and multivariate metrics.

**Figure S2:**
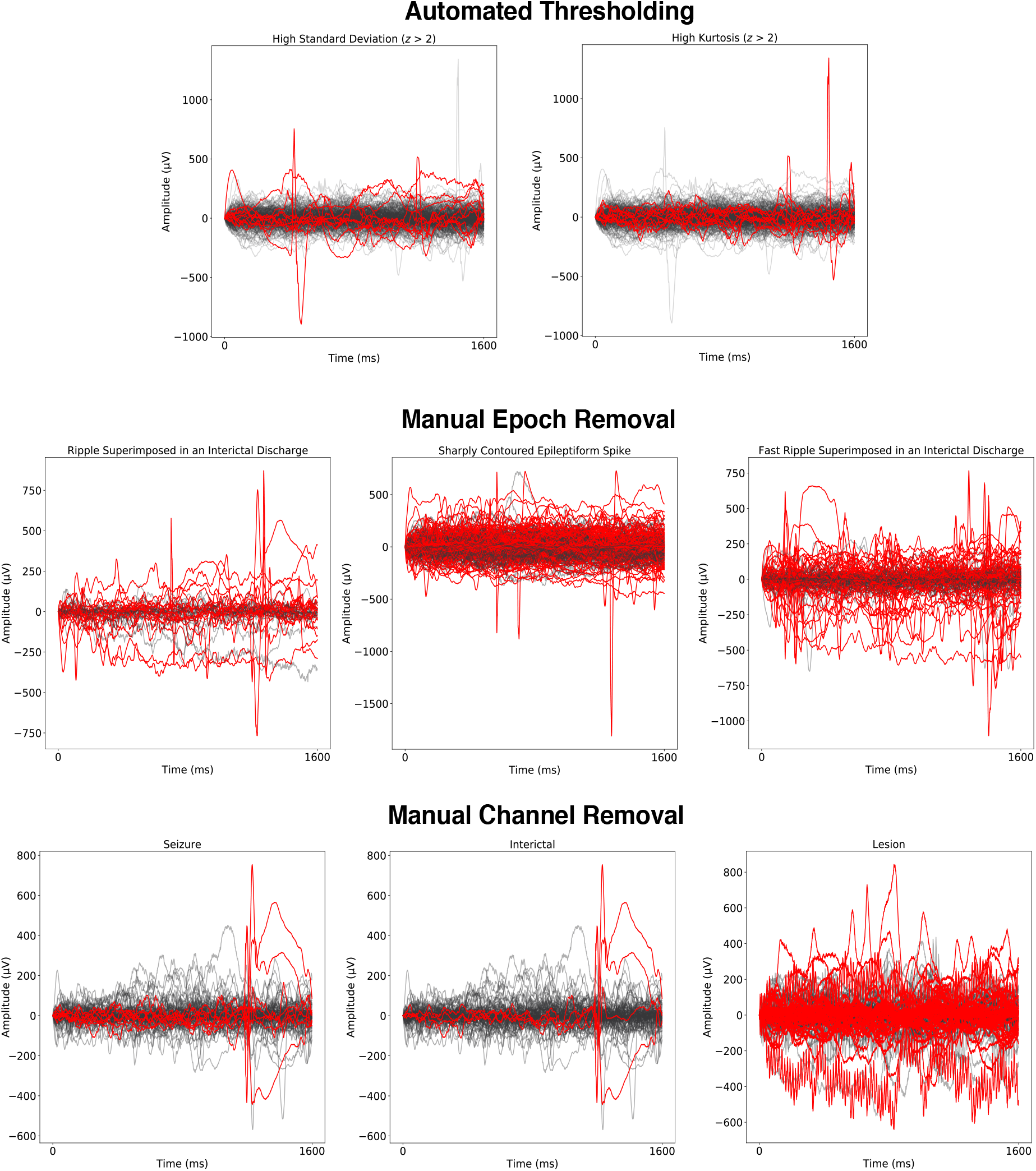
Examples of timeseries that each data cleaning method excluded. Red lines represents data from a removed epoch or channel, while black segments represent data retained by the method. All depicted signals have been bipolar referenced and represent an epoch’s worth of information (1600 ms) from a single subject. Each bold heading denotes a different method, with individual plot titles describing the reason for exclusion.

**Figure S3:**
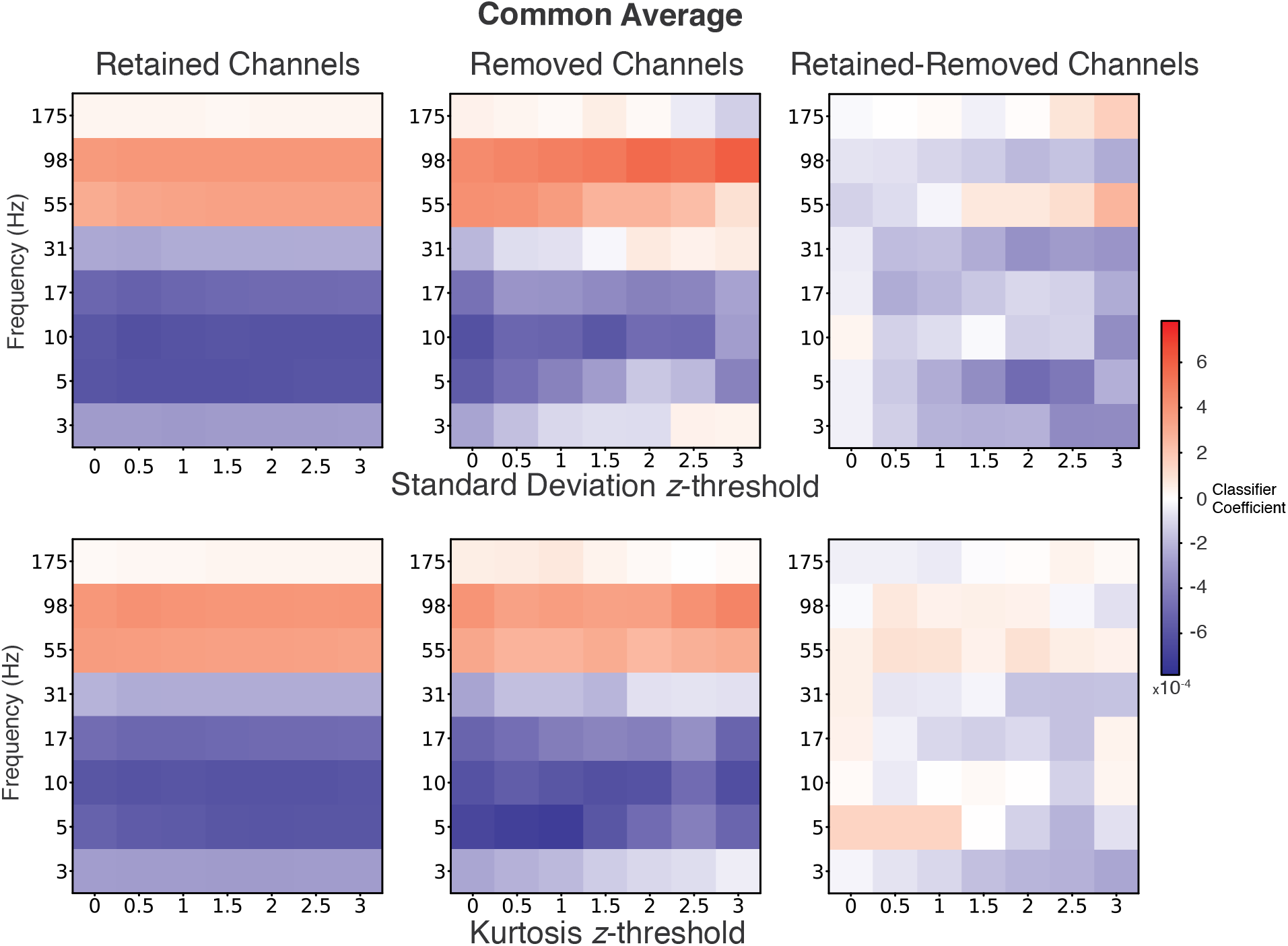
Average classifier coefficients in common-average referenced data for electrodes retained and excluded at the tested thresholds, as well as the difference between the two sets of coefficients. No significant differences after FDR correction (*q* < 0.05).

**Figure S4:**
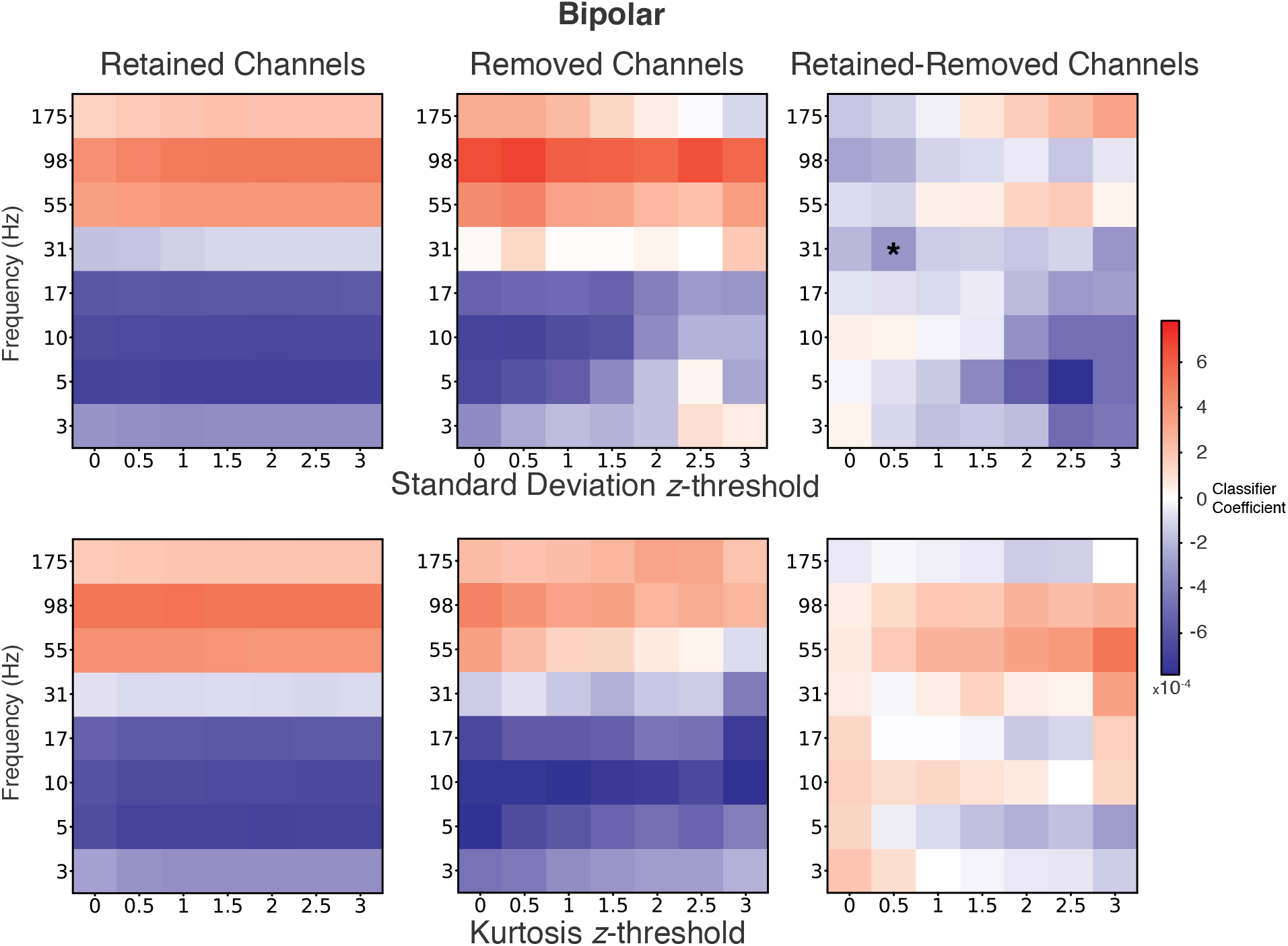
Average classifier coefficients in bipolar referenced data for electrodes retained and excluded at the tested thresholds, as well as the difference between the two sets of coefficients. * Denotes significant difference between excluded and retained following FDR correction (Wilcoxon signed-rank test, *q* < 0.05).

**Figure S5:**
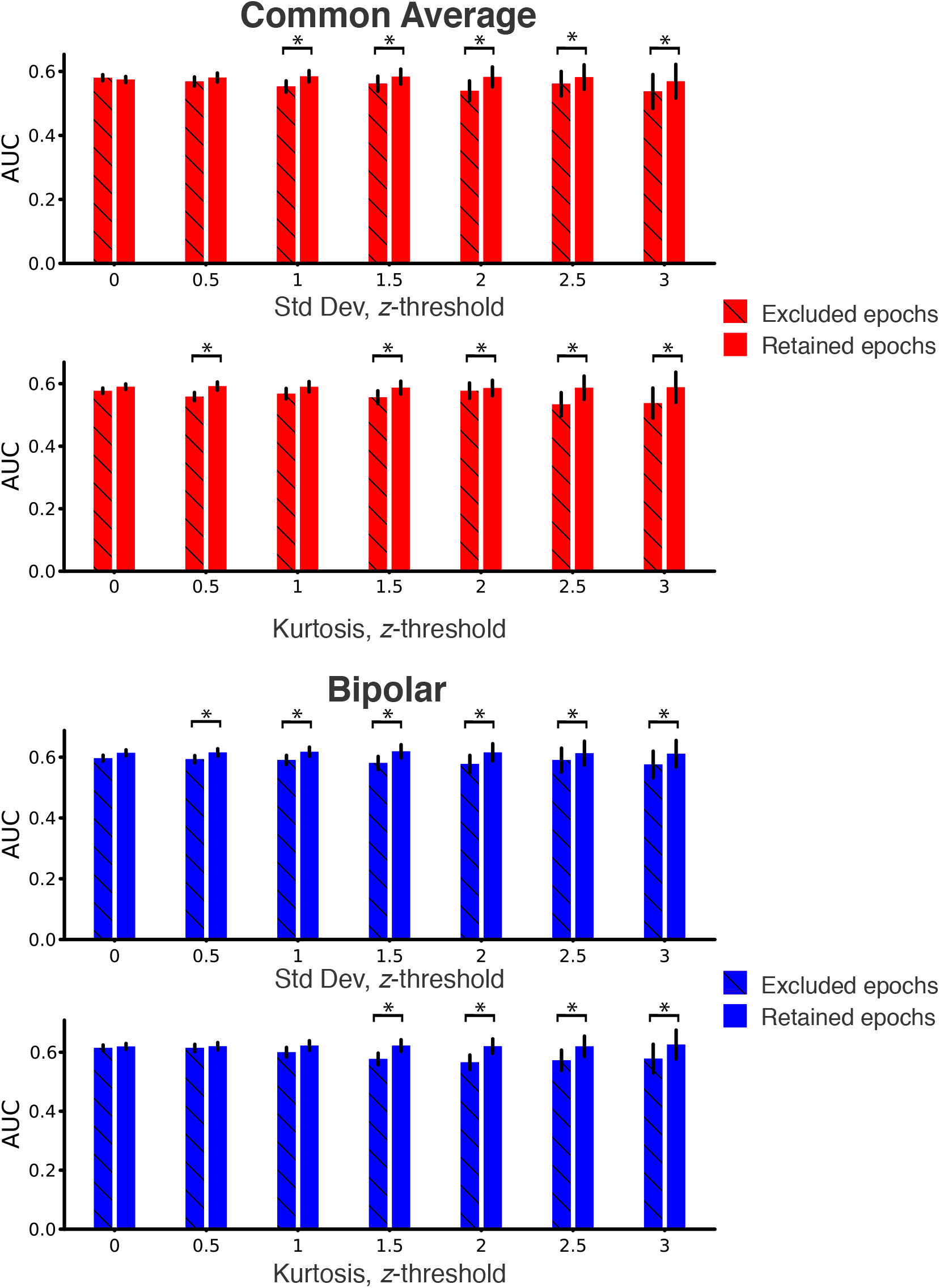
Comparison of how accurately excluded and retained epochs are classified correctly at the tested exclusion thresholds. The recall probabilities for each epoch was derived from a classifier trained without excluding any data. Error bars represent SEM. * Denotes significant difference following FDR correction (Wilcoxon signed-rank test, *q* < 0.05).

**Figure S6:**
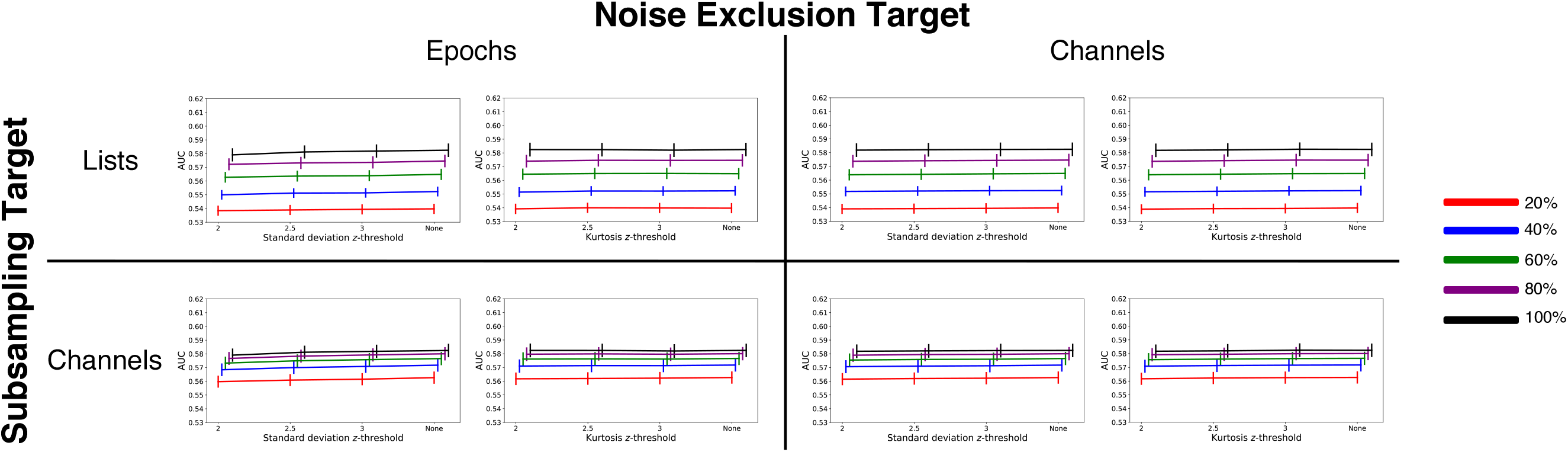
Multivariate results after subsampling common average rereferenced data. Different colors denote different percentages of subsampled data. Red: 20%, Blue: 40%, Green: 60%, Purple: 80%, Black: 100% (no subsampling). Error bars represent SEM. Subsampling to lower deciles of data (both in term of number of events and channels) did not lead to significant interactions with noise removal.

**Figure S7:**
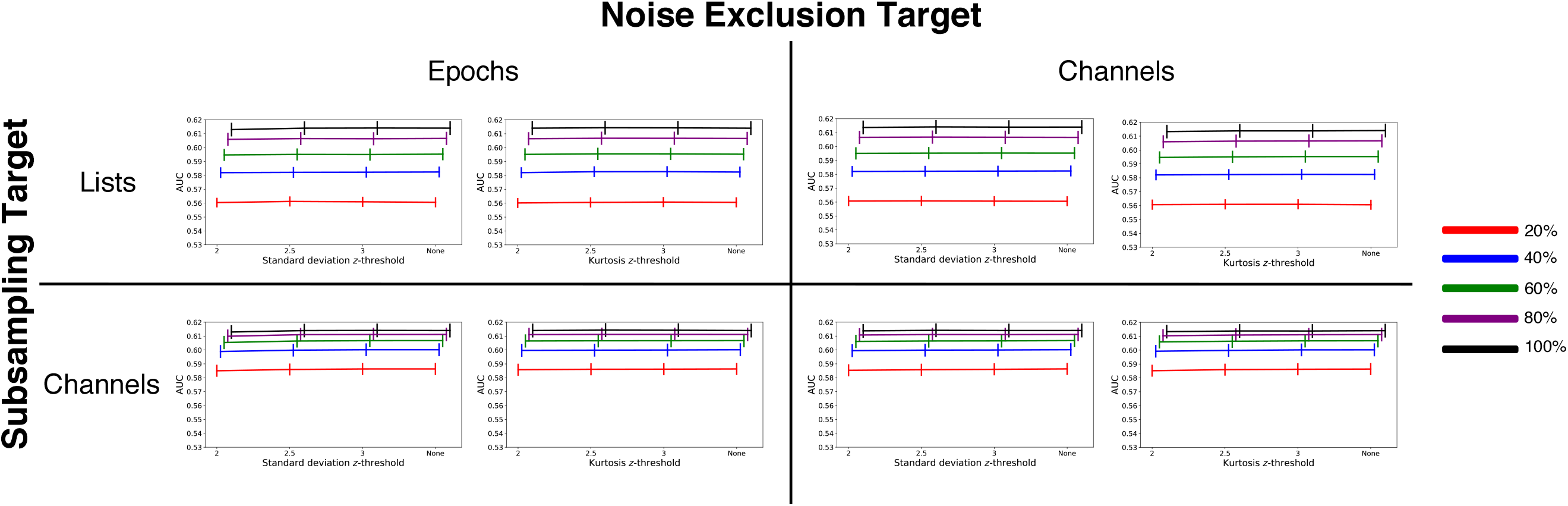
Multivariate results after subsampling bipolar rereferenced data. Different colors denote different percentages of subsampled data. Red: 20%, Blue: 40%, Green: 60%, Purple: 80%, Black: 100% (no subsampling). Error bars represent SEM. Subsampling to lower deciles of data (both in term of number of events and channels) did not lead to significant interactions with noise removal.

**Figure S8:**
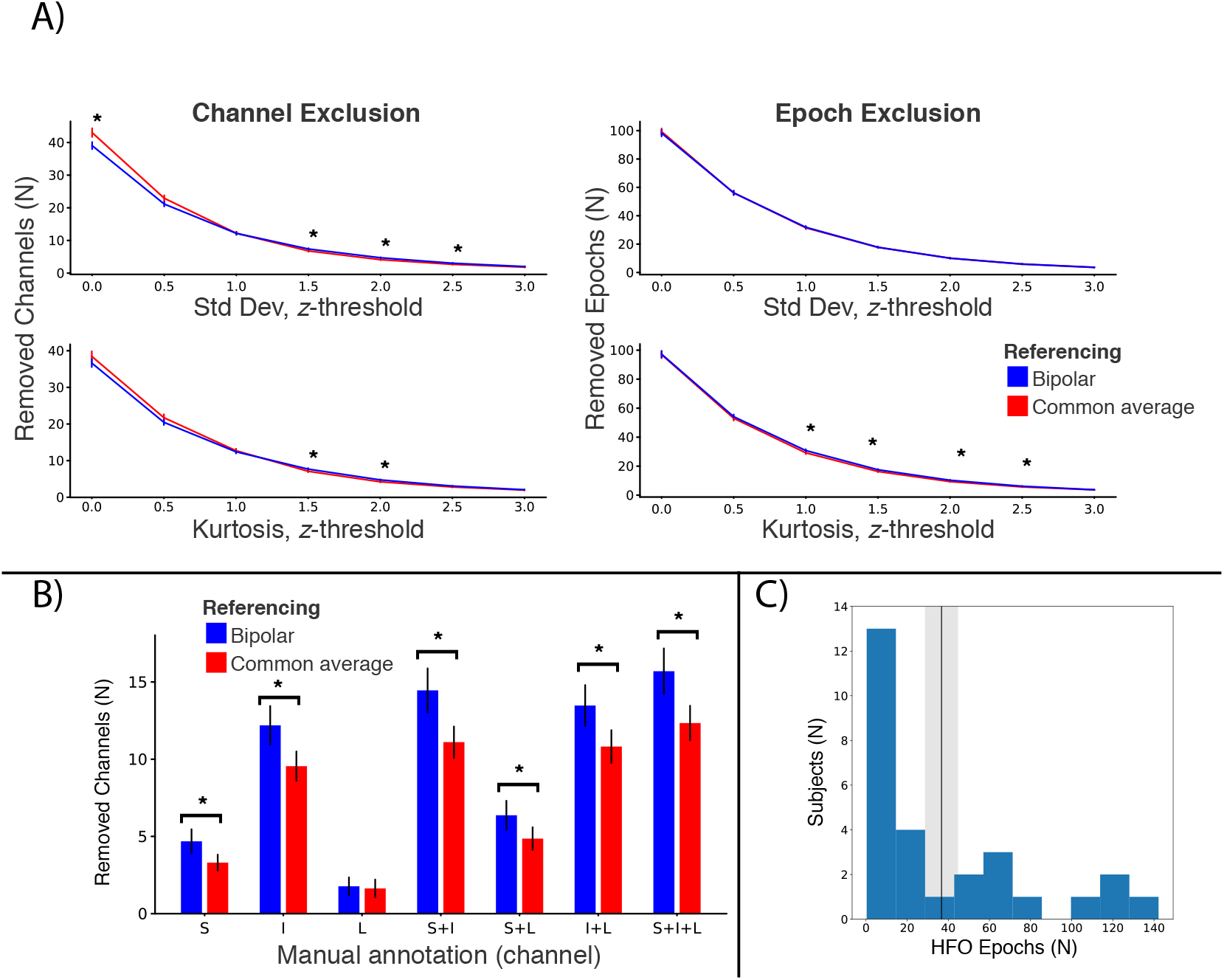
Depiction of how much data was removed by each method. Error bars represent SEM. A) Amount of epochs and channels removed by automated thresholding. B) Number of expert-reviewed abnormal annotated channels. S: Seizure, I: Interictal, L: Lesion. C) Number of HFO epochs in our subject cohort. The black line and shaded region surrounding it denote the average and SEM, respectively.

## Notes

http://memory.psych.upenn.edu/Electrophysiological_Data

## References

Bootkrajang, J., & Kabán, A. (2012). Label-noise robust logistic regression and its applications. In Joint european conference on machine learning and knowledge discovery in databases (pp. 143–158).

Burke, J. F., Long, N. M., Zaghloul, K. A., Sharan, A. D., Sperling, M. R., & Kahana, M. J. (2014). Human intracranial high-frequency activity maps episodic memory formation in space and time. NeuroImage, 85, 834–843.

Burke, J. F., Zaghloul, K. A., Jacobs, J., Williams, R. B., Sperling, M. R., Sharan, A. D., & Kahana, M. J. (2013). Synchronous and asynchronous theta and gamma activity during episodic memory formation. Journal of Neuroscience, 33(1), 292–304.

Caplan, J. B., Madsen, J. R., Raghavachari, S., & Kahana, M. J. (2001). Distinct patterns of brain oscillations underlie two basic parameters of human maze learning. Journal of Neurophysiology, 86, 368–380.

Carroll, R. J., & Pederson, S. (1993). On robustness in the logistic regression model. Journal of the Royal Statistical Society. Series B (Methodological), 693–706.

Chang, E. F., Edwards, E., Nagarajan, S. S., Fogelson, N., Dalal, S. S., Canolty, R. T., … Knight, R. T. (2011). Cortical spatio-temporal dynamics underlying phonological target detection in humans. Journal of cognitive neuroscience, 23(6), 1437–1446.

Cheyne, D., Bells, S., Ferrari, P., Gaetz, W., & Bostan, A. C. (2008). Self-paced movements induce high-frequency gamma oscillations in primary motor cortex. Neuroimage, 42(1), 332–342.

Crone, N. E., Sinai, A., & Korzeniewska, A. (2006). High-frequency gamma oscillations and human brain mapping with electrocorticography. Progress in Brain Research, 159, 275–295.

Dastjerdi, M., Foster, B. L., Nasrullah, S., Rauschecker, A. M., Dougherty, R. F., Townsend, J. D., … others (2011). Differential electrophysiological response during rest, self-referential, and non-self-referential tasks in human posteromedial cortex. Proceedings of the National Academy of Sciences, 108(7), 3023–3028.

Ekstrom, A. D., Caplan, J., Ho, E., Shattuck, K., Fried, I., & Kahana, M. (2005). Human hippocampal theta activity during virtual navigation. Hippocampus, 15, 881–889.

Ezzyat, Y., Kragel, J. E., Burke, J. F., Levy, D. F., Lyalenko, A., Wanda, P.,… Kahana, M. J. (2017). Direct brain stimulation modulates encoding states and memory performance in humans. Current Biology, 27(9), 1251–1258.

Ezzyat, Y., & Rizzuto, D. S. (2018). Direct brain stimulation during episodic memory. Current Opinion in Biomedical Engineering, 8, 78–83.

Ezzyat, Y., Wanda, P., Levy, D. F., Kadel, A., Aka, A., Pedisich, I., … Kahana, M. J. (2018). Closed-loop stimulation of temporal cortex rescues functional networks and improves memory. Nature Communications, 9(1), 365. doi: 10.1038/s41467-017-02753-0

Fonken, Y. M., Rieger, J. W., Tzvi, E., Crone, N. E., Chang, E., Parvizi, J., … Krämer, U. M. (2016). Frontal and motor cortex contributions to response inhibition: evidence from electrocorticography. Journal of neurophysiology, 115(4), 2224–2236.

Foster, B. L., Kaveh, A., Dastjerdi, M., Miller, K. J., & Parvizi, J. (2013). Human retrosplenial cortex displays transient theta phase locking with medial temporal cortex prior to activation during autobiographical memory retrieval. The Journal of Neuroscience, 33(25), 10439–10446.

García-Cordero, I., Esteves, S., Mikulan, E. P., Hesse, E., Baglivo, F. H., Silva, W., … others (2017). Attention, in and out: scalp-level and intracranial eeg correlates of interoception and exteroception. Frontiers in neuroscience, 11, 411.

Glanzer, M. (1969). Distance between related words in free recall: Trace of the STS. Journal of Verbal Learning and Verbal Behavior, 8, 105–111.

Greco, A., Mammone, N., Morabito, F., & Versaci, M. (2007). Semi-automatic artifact rejection procedure based on kurtosis, Renyi’s entropy and independent component scalp maps. International Journal of Medical, Health, Biomedical and Pharmaceutical Engeneering, 1, 466–470.

Greenberg, J. A., Burke, J. F., Haque, R., Sharan, A. D., Litt, B., Baltuch, G. H.,… Zaghloul, K. A. (2015). Decreases in theta and increases in high frequency activity underlie associative memory encoding. NeuroImage, 114, 257–263.

Hammon, P. S., & de Sa, V. R. (2007). Preprocessing and meta-classification for brain-computer interfaces. IEEE Transactions on Biomedical Engineering, 54(3), 518–525.

Hanley, J. A., & McNeil, B. J. (1982). The meaning and use of the area under a receiver operating characteristic (roc) curve. Radiology, 143(1), 29–36.

Haque, R. U., Wittig, J. H., Damera, S. R., Inati, S. K., & Zaghloul, K. A. (2015). Cortical low-frequency power and progressive phase synchrony precede successful memory encoding. The Journal of Neuroscience, 35(40), 13577–13586.

Hastie, T., Tibshirani, R., & Friedman, J. (2001). The elements of statistical learning. New York: Springer-Verlag.

Hermes, D., Miller, K. J., Wandell, B. A., & Winawer, J. (2015). Gamma oscillations in visual cortex: the stimulus matters. Trends in cognitive sciences, 19(2), 57–58.

Horak, P., Meisenhelter, S., Song, Y., Testorf, M., Kahana, M., Viles, W. D., … Jobst, B. C. (2017). Interictal epileptiform discharges impair word recall in multiple brain areas. Epilepsia, 58(3), 373–380.

Kim, K., Ekstrom, A. D., & Tandon, N. (2016). A network approach for modulating memory processes via direct and indirect brain stimulation: Toward a causal approach for the neural basis of memory. Neurobiology of Learning and Memory.

Komarek, P., & Moore, A. W. (2003). Fast robust logistic regression for large sparse datasets with binary outputs. In Aistats.

Lachaux, J. P., Axmacher, N., Mormann, F., Halgren, E., & Crone, N. E. (2012). High-frequency neural activity and human cognition: Past, present, and possible future of intracranial EEG research. Progress in Neurobiology, 98, 279–301.

Lega, B., Burke, J. F., Jacobs, J., & Kahana, M. J. (2015). Slow theta-to-gamma phase amplitude coupling in human hippocampus supports the formation of new episodic memories. Cerebral Cortex, 26(1), 268–278.

Lega, B., Kahana, M. J., Jaggi, J. L., Baltuch, G. H., & Zaghloul, K. A. (2011). Neuronal and oscillatory activity during reward processing in the human ventral striatum. NeuroReport, 22(16), 795–800.

Li, G., Jiang, S., Paraskevopoulou, S. E., Wang, M., Xu, Y., Wu, Z., … Schalk, G. (2018). Optimal referencing for stereo-electroencephalographic (seeg) recordings. NeuroImage, 183, 327–335.

Long, N. M., Burke, J. F., & Kahana, M. J. (2014). Subsequent memory effect in intracranial and scalp EEG. NeuroImage, 84, 488–494. doi: 10.1016/j.neuroimage.2013.08.052

Long, N. M., & Kahana, M. J. (2015). Successful memory formation is driven by contextual encoding in the core memory network. NeuroImage, 119, 332–337.

Manning, J. R., Jacobs, J., Fried, I., & Kahana, M. J. (2009). Broadband shifts in local field potential power spectra are correlated with single-neuron spiking in humans. Journal of Neuroscience, 29(43), 13613–13620.

Mercier, M. R., Bickel, S., Megevand, P., Groppe, D. M., Schroeder, C. E., Mehta, A. D., & Lado, F. A. (2017). Evaluation of cortical local field potential diffusion in stereotactic electro-encephalography recordings: a glimpse on white matter signal. Neuroimage, 147, 219–232.

Michelmann, S., Treder, M. S., Griffiths, B., Kerren, C., Roux, F., Wimber, M., … others (2018). Data-driven re-referencing of intracranial eeg based on independent component analysis (ica). Journal of neuroscience methods, 307, 125–137.

Miller, K., den Nijs, M., Shenoy, P., Miller, J., Rao, R., & Ojemann, J. (2007). Real-time functional brain mapping using electrocorticography. Neuroimage, 37(2), 504–507.

Miller, K. J., Leuthardt, E. C., Schalk, G., Rao, R. P. N., Anderson, N. R., Moran, D. W., … Ojemann, J. G. (2007). Spectral changes in cortical surface potentials during motor movement. Journal of Neuroscience, 27, 2424–2432.

Nikulin, V. V., Nolte, G., & Curio, G. (2011). A novel method for reliable and fast extraction of neuronal eeg/meg oscillations on the basis of spatio-spectral decomposition. NeuroImage, 55(4), 1528–1535.

Nolan, H., Whelan, R., & Reilly, R. (2010). Faster: fully automated statistical thresholding for eeg artifact rejection. Journal of neuroscience methods, 192(1), 152–162.

Nunez, P. L., & Srinivasan, R. (2006). Electric fields of the brain. New York: Oxford University Press.

Parvizi, J., & Kastner, S. (2018). Promises and limitations of human intracranial electroencephalography. Nature neuroscience, 1.

Pedregosa, F., Varoquaux, G., Gramfort, A., Michel, V., Thirion, B., Grisel, O., … Duchesnay, E. (2011). Scikit-learn: Machine learning in Python. Journal of Machine Learning Research, 12, 2825–2830.

Penny, W. D., Roberts, S. J., Curran, E. A., & Stokes, M. J. (2000). Eeg-based communication: a pattern recognition approach. IEEE transactions on Rehabilitation Engineering, 8(2), 214–215.

Phan, T. D., Wachter, J. A., & Kahana, M. J. (Submitted). Multivariate stochastic volatility modeling of neural data. Submitted.

Piai, V., Anderson, K. L., Lin, J. J., Dewar, C., Parvizi, J., Dronkers, N. F., & Knight, R. T. (2016). Direct brain recordings reveal hippocampal rhythm underpinnings of language processing. Proceedings of the National Academy of Sciences, 113(40), 11366–11371.

Raghavachari, S., Kahana, M. J., Rizzuto, D. S., Caplan, J. B., Kirschen, M. P., Bourgeois, B., … Lisman, J. E. (2001). Gating of human theta oscillations by a working memory task. Journal of Neuroscience, 21(9), 3175–3183.

Rangarajan, V., Hermes, D., Foster, B. L., Weiner, K. S., Jacques, C., Grill-Spector, K., & Parvizi, J. (2014). Electrical stimulation of the left and right human fusiform gyrus causes different effects in conscious face perception. Journal of Neuroscience, 34(38), 12828–12836.

Rutishauser, U., Ross, I., Mamelak, A., & Schuman, E. (2010). Human memory strength is predicted by theta-frequency phase-locking of single neurons. Nature, 464(7290), 903–907.

Ryali, S., Supekar, K., Abrams, D. A., & Menon, V. (2010). Sparse logistic regression for whole-brain classification of fmri data. NeuroImage, 51(2), 752–764.

Sederberg, P. B., Gauthier, L. V., Terushkin, V., Miller, J. F., Barnathan, J. A., & Kahana, M. J. (2006). Oscillatory correlates of the primacy effect in episodic memory. NeuroImage, 32(3), 1422–1431. doi: 10.1016/j.neuroimage.2006.04.223

Sederberg, P. B., Kahana, M. J., Howard, M. W., Donner, E. J., & Madsen, J. R. (2003). Theta and gamma oscillations during encoding predict subsequent recall. Journal of Neuroscience, 23(34), 10809–10814.

Sederberg, P. B., Schulze-Bonhage, A., Madsen, J. R., Bromfield, E. B., Litt, B., Brandt, A., & Kahana, M. J. (2007). Gamma oscillations distinguish true from false memories. Psychological Science, 18(11), 927–932.

Sheehan, T. C., Sreekumar, V., Inati, S. K., & Zaghloul, K. A. (2018). Signal complexity of human intracranial eeg tracks successful associative memory formation across individuals. Journal of Neuroscience, 2389–17.

Shimamoto, S., Waldman, Z. J., Orosz, I., Song, I., Bragin, A., Fried, I.,… others (2018). Utilization of independent component analysis for accurate pathological ripple detection in intracranial eeg recordings recorded extra-and intra-operatively. Clinical Neurophysiology, 129(1), 296–307.

Silberzahn, R., Uhlmann, E. L., Martin, D. P., Anselmi, P., Aust, F., Awtrey, E., … others (2018). Many analysts, one data set: Making transparent how variations in analytic choices affect results. Advances in Methods and Practices in Psychological Science, 1(3), 337–356.

Simmons, J. P., Nelson, L. D., & Simonsohn, U. (2011). False-positive psychology: Undisclosed flexibility in data collection and analysis allows presenting anything as significant. Psychological science, 22(11), 1359–1366.

Solomon, E., Kragel, J., Sperling, M., Sharan, A., Worrell, G., Kucewicz, M., … Kahana, M. (2017). Widespread theta synchrony and high-frequency desynchronization underlies enhanced cognition. Nature Communications, 8(1), 1704. doi: 10.1038/s41467-017-01763-2

Swann, N. C., Tandon, N., Pieters, T. A., & Aron, A. R. (2012). Intracranial electroencephalography reveals different temporal profiles for dorsal-and ventro-lateral prefrontal cortex in preparing to stop action. Cerebral Cortex, 23(10), 2479–2488.

Tomioka, R., Aihara, K., & Müller, K.-R. (2007). Logistic regression for single trial eeg classification. In Advances in neural information processing systems (pp. 1377–1384).

van Vugt, M. K., Schulze-Bonhage, A., Litt, B., Brandt, A., & Kahana, M. J. (2010). Hippocampal gamma oscillations increase with memory load. Journal of Neuroscience, 30(7), 2694–2699.

van Vugt, M. K., Schulze-Bonhage, A., Sekuler, R., Litt, B., Brandt, A., Baltuch, G., & Kahana, M. J. (2009). Intracranial electroencephalography reveals two distinct similarity effects during item recognition. Brain Research, 1299, 33–44. doi: 10.1016/j.brainres.2009.07.016

Vass, L. K., Copara, M. S., Seyal, M., Shahlaie, K., Farias, S. T., Shen, P. Y., & Ekstrom, A. D. (2016). Oscillations go the distance: Low-frequency human hippocampal oscillations code spatial distance in the absence of sensory cues during teleportation. Neuron, 89(6), 1180–1186.

Voytek, B., Canolty, R., Shestyuk, A., Crone, N., Parvizi, J., & Knight, R. (2010). Shifts in gamma phase–amplitude coupling frequency from theta to alpha over posterior cortex during visual tasks. Frontiers in Human Neuroscience, 4.

Waldman, Z. J., Shimamoto, S., Song, I., Orosz, I., Bragin, A., Fried, I.,… Weiss, S. A. (2018). A method for the topographical identification and quantification of high frequency oscillations in intracranial electroencephalography recordings. Clinical Neurophysiology, 129(1), 308–318.

Weiss, S. A., Berry, B., Chervoneva, I., Waldman, Z., Guba, J., Bower, M., … others (2018). Visually validated semi-automatic high-frequency oscillation detection aides the delineation of epileptogenic regions during intra-operative electrocorticography. Clinical Neurophysiology, 129(10), 2089–2098.

Whitmer, D., Worrell, G., Stead, M., Lee, I. K., & Makeig, S. (2010). Utility of independent component analysis for interpretation of intracranial eeg. Frontiers in human neuroscience, 4, 184.

Winawer, J., Kay, K. N., Foster, B. L., Rauschecker, A. M., Parvizi, J., & Wandell, B. A. (2013). Asynchronous broadband signals are the principal source of the bold response in human visual cortex. Current Biology, 23(13), 1145–1153.

Zavala, B. A., Tan, H., Little, S., Ashkan, K., Hariz, M., Foltynie, T., … Brown, P. (2014). Midline frontal cortex low-frequency activity drives subthalamic nucleus oscillations during conflict. Journal of Neuroscience, 34(21), 7322–7333.

Zhang, H., & Jacobs, J. (2015). Traveling theta waves in the human hippocampus. The Journal of Neuroscience, 35(36), 12477–12487.

